# Chloroplast genome of the nutmeg tree: Myristica fragrans Houtt. (Myristicaceae)

**DOI:** 10.1101/2020.02.25.964122

**Authors:** Sylvia Mota de Oliveira, Elza Duijm, Hans ter Steege

## Abstract

*Myristica fragrans* Houtt. (Myristicaceae) is widely used as condiment in western countries as well as a drug in medicinal systems such as the Ayuverda and Unani. The assemblage of its chloroplast genome resulted in a total of 155,894 bp, from which 146 genes were annotated, along with 86 coding regions, 43 exons and 22 introns. This study is a step further in the species characterization and will support future phylogenetic studies within the family.

## Introduction

*Myristica fragrans* Houtt. is the most important species of the plant family Myristicaceae in the global market. The tree bears fruits containing oblong seeds, wrapped in a red aril. The world export volume of these seeds and arils, namely nutmeg and mace, attained a peak of 15,501 tons in 2011 (http://www.fao.org/). The regulation of the international trade is mainly concerned with the use of these products in western countries as spices, and therefore guarantees the presence of its principal constituents, such as steam volatile oil (essential oil), fixed (fatty) oil, proteins, cellulose, pentosans, starch, resin and mineral elements (http://www.fao.org/3/x5047E/x5047E0a.htm). However, there is an even stronger demand for good characterization of *M. fragrans* as highly important medicinal plant available in the market (Mishra et al. 2016). The dried arils of the seeds are used since ancient times in drug preparation, being part of different medicinal systems, such as the Ayuverda and Unani. Myristicin, the active principle of the drug, has been the focus of several pharmacological studies and its isolation, characterization and quantification, aimed to develop standard parameters for the genuine drug (Naikodi et al. 2011). Experimental evidence for the use of *M. argentea* and *M. malabarica* as substitutes/adulterants of *M. fragrans* in medicinal herbal market demonstrates the need of good species identification even when only plant fragments are available (Dhanya and Sasikumar 2010). In this line, molecular data has been also incorporated as a tool for herbal product authentication (Swethaab et al. 2017).

The use of molecular data as a species barcode in plants went through several discussions since the first attempts to select standard DNA regions (Hollingsworth et al. 2009, Kress et al. 2009). In the last decades, sequences of the chloroplast regions matK and rbcL, together with nuclear ITS, have become available for a large number of plants and extensively used in phylogenies. But the limited amount of information in these selected regions may lead to unresolved phylogenies, depending on the taxon (Hollingsworth et al. 2016). From previous studies with selected regions, it is known that there is remarkably low amount of molecular variation in Myristicaceae, which makes it difficult to reconstruct the relationship among its genera (Sauquet et al. 2003).

The latest developments of high throughput sequence technology platforms as well as dedicated software for data analysis, coupled with the reduction of data generation costs, stimulated the field towards whole genome data. Through this approach, the suggested barcode regions can be recovered, but the complete plastid genome becomes available as well. The assemblage and annotation of the chloroplast genome of *Myristica fragrans* presented here is a further step towards species characterization as well as a reference for future phylogenetic studies on Myristicaceae.

## Material and Methods

Fresh leaf samples were taken from one individual of *Myristica fragrans* growing in the Hortus Botanicus Leiden (The Netherlands). Enrichment of the chloroplast from the leaf samples was performed using ± 2 cm^2^ of leaf tissue with the Chloroplast Isolation Kit (Sigma, CPISO-1KT After this the DNA was extracted using a modified C-Tab protocol (Doyle JJ, Doyle JL 1987). Concentration was determined using the Dropsense and the quality of the DNA was checked on the QIAxcel (QIAgen). Sonification was performed with the Covaris M220 (covaris), using the microTUBE-59 AFA Fiber Screw-Cap according to the manufacturer’s program for an insert size of 350 bp. Followed by a Dual index library prep with the DNA NEBNext^®^ Ultra™ II Library Prep Kit (New England Biolabs), using a quarter of the recommended volumes with the Dual Index Primers Set, NEBNext^®^ Multiplex Oligos for Illumina^®^. The libraries were checked for concentration with the QIAxcel (Qiagen) and pooled using the QIAgility (QIAgen). A final quantity and quality check was performed with the Bioanalyzer (Agilent) using a High Sensitivity chip. Paired-end sequence reads of 150 bp were generated using the Illumina NovaSeq 6000 system.

The sequences generated with the NovaSeq system were performed under accreditation according to the scope of BaseClear B.V. (L457; NEN-EN-ISO/IEC 17025). When paired-end sequencing is being performed, “number of reads” refers to read pairs. FASTQ read sequence files were generated using bcl2fastq2 version 2.18. Initial quality assessment was based on data passing the Illumina Chastity filtering. Subsequently, reads containing PhiX control signal were removed using an in-house filtering protocol. In addition, reads containing (partial) adapters were clipped (up to a minimum read length of 50 bp). The second quality assessment was based on the remaining reads using the FASTQC quality control tool version 0.11.5.

The chloroplast genome was preliminary assembled using two *denovo* assembly tools, NOVOPlasty (Dierckxsens et al 2017) and GetOrganelle (Jian-Jun et al 2019), to explore the consistency of the results. Further assembly was carried out with NOVOPlasty, using the rbcL of *M. fragrans* (NCBI accession MH069804.1) as seed. Genome annotation was carried out with GeSeq Version 1.76 (Tilich et al 2017) using the chloroplast genome of *Horsfieldia pandurifolia* as reference sequence (Mao et al 2019). Then we constructed a visual representation of the plastome using OGDRAW (Greiner et al 2019). We used Minimap2 (Li 2018) and Samtools (Li et al 2009) to map all chloroplast reads of the original paired end Illumina files on the de-novo assembled plastome, to check for consistency and depth of coverage.

Finally, we used published chloroplast sequences deposited in NCBI of another four species of Myristicaceae viz. *Knema elegans, Knema furfuracea, Myristica yunnanensis, Horsfieldia pandurifolia* (NCBI accession respectively: MK285564.1, MK285563.1, MK285565, NC_042225.1) and two of Magnoliaceae viz. *Michelia alba* and *Liriodendron tulipifera* (NCBI accession respectively: NC_037005.1; DQ899947.1), to construct a distance tree including the data of our *Myristica fragrans* assembled genome. The distance tree was built in R, using libraries “kmer”, “ape” and “vegan” with kmer size 10 and Jaccard distance.

## Results

### Sequencing

Nucleotides statistics are given as supplemental online material (S1). The Illumina products corresponded to 13,224,370 sequences, 95% of which had a length of 151 base pairs (bp) and the remaining 5% increasingly approached 150 bp (S1; Fig. 1). Minimum length was 50 bp. Frequency of A, C, G and T bases were 30%, 20.1%, 20.3% and 29.6%. Records solved as N were negligible (<1%). GC content reached 40.4%.

The mapping of the reads on the assembled genome showed relatively stable coverage across the genome (S1; Fig. 2), with an average depth of 884 copies. The exceptions, where coverage dropped to less than 500 copies, were in the positions corresponding to part of rps19 fragment gene and trnH-GUG gene, the non-annotated region between rpl20 and rps12 (around position 72296 in our data), and the region before trnA-UGC.

### *Denovo* assemblage and annotation

After assemblage, the two samples of fresh material delivered the same number of contigs and total length. Due to the consistency of the results, we picked the best performing sample in terms of number of reads – sn2 – to produce the final assemblage. In both NOVOPlasty and GetOrganelle, the assembled genome had a total length of 155,894 bp distributed in three contigs that could be concatenated with two different scaffolds. We used the assembly performed by NOVOPlasty as final result. It generated 3 contigs with lengths of 135,207 bp, 21,808 bp and 21,715 bp with an average insert size of 265 bp.

Organelle genome corresponded to 3.86% of the bulk material. The differences between the two scaffolds consisted of differences in the orientation, forward or reverse, of the following regions: ccsA, ndhA, ndhD, ndhE, ndhF, ndhG, ndhH, ndhl, psaC, rpl32, rps15, trnL-UAG, trnN-GUU, trnR-ACG. The comparison of gene orientation between each of the options against the complete plastome of *Horsfieldia pandurifolia* guided our choice of scaffold.

The annotation of the chloroplast genome of *Myristica fragrans* corresponded to the expected pattern of one large single copy and one small single copy separated by two inverted repeat regions. A total of 146 genes were annotated, along with 86 coding regions, 43 exons and 22 introns (Fig. 1). Molecular data is available at NCBI platform under NCBI accession MT062914.

The distance tree based on kmer analysis including *M. fragrans*, another four species of Myristicaceae and two Magnoliaceae increased the reliability of the genome assemblage of *M. fragrans* because it recovered the expected relationships among taxa (Fig. 2).

The high coverage of the chloroplast genome of *Myristica fragrans* allowed *denovo* assembly with high reliability. Only two scaffold options were possible. Our choice was based on the orientation of 14 regions, compared to previously published Myristicaceae species. The robustness of the results is also supported by the fact that both assemblage tools used, NOVOPlasty (Dierckxsens et al 2017) and GetOrganelle (Jian-Jun et al 2019), provided the same solution for our dataset.

## Supporting information

Supplemental nucleotide statistics

## Acknowledgments

We thank the Hortus Botanicus Leiden and its staff, especially Guilherme Herrero Bastos, for growing *Myristica fragrans* in their greenhouse and providing us with leaf samples.

---

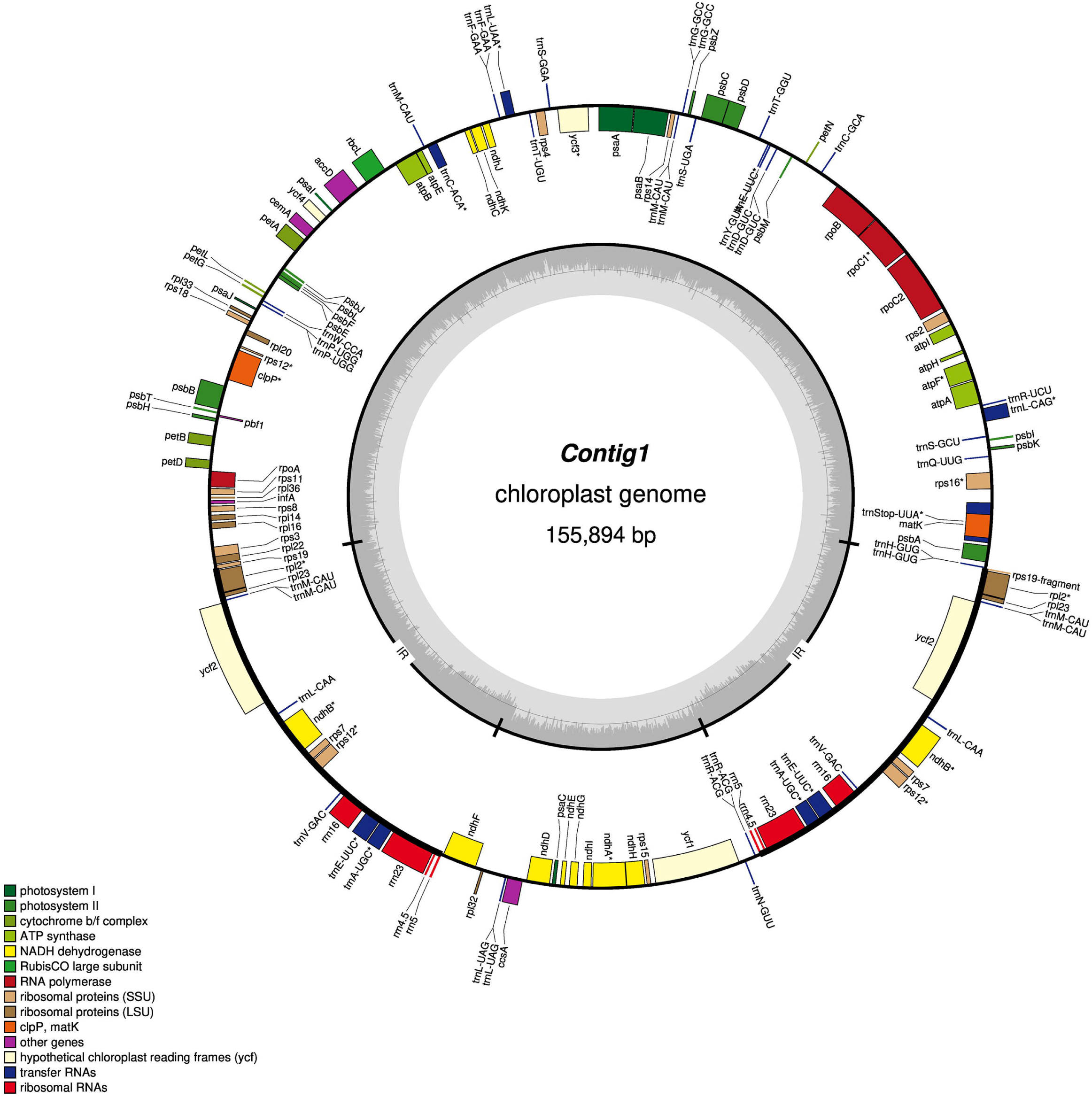

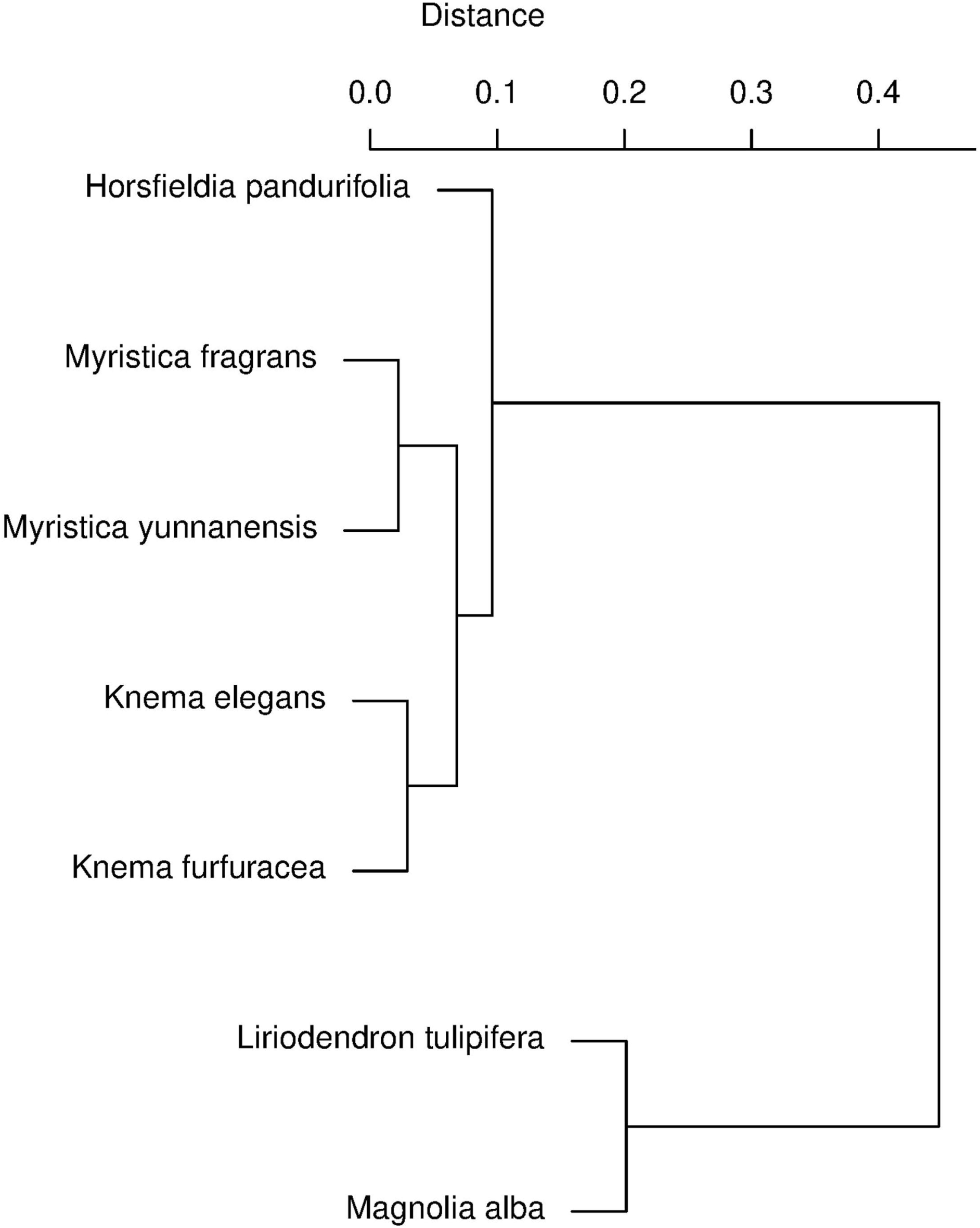

