## Supplemental nucleotide statistics for "Chloroplast genome of the nutmeg tree: Myristica fragrans Houtt. (Myristicaceae)"

Length: 151 bp

Sequences: 13,224,370

Lengths: Mean: 149.6; Std Dev: 8.7; Minimum: 50; Maximum: 151

Confidence Mean: 35.7

Expected Errors: 5,215,652

Error Free Odds: < 0.0001%

At least Q20: 97.1%

At least Q30: 92.1%

At least Q40: 0.0%

Table 1. Frequency bases:

| Base | Absolute | Percentage |
| --- | --- | --- |
| A | 593,282,422 | 30.0% |
| C | 397,988,842 | 20.1% |
| G | 400,898,179 | 20.3% |
| T | 585,830,256 | 29.6% |
| N | 14,426 | 0.0% |
| GC | 798,887,021 | 40.4% |

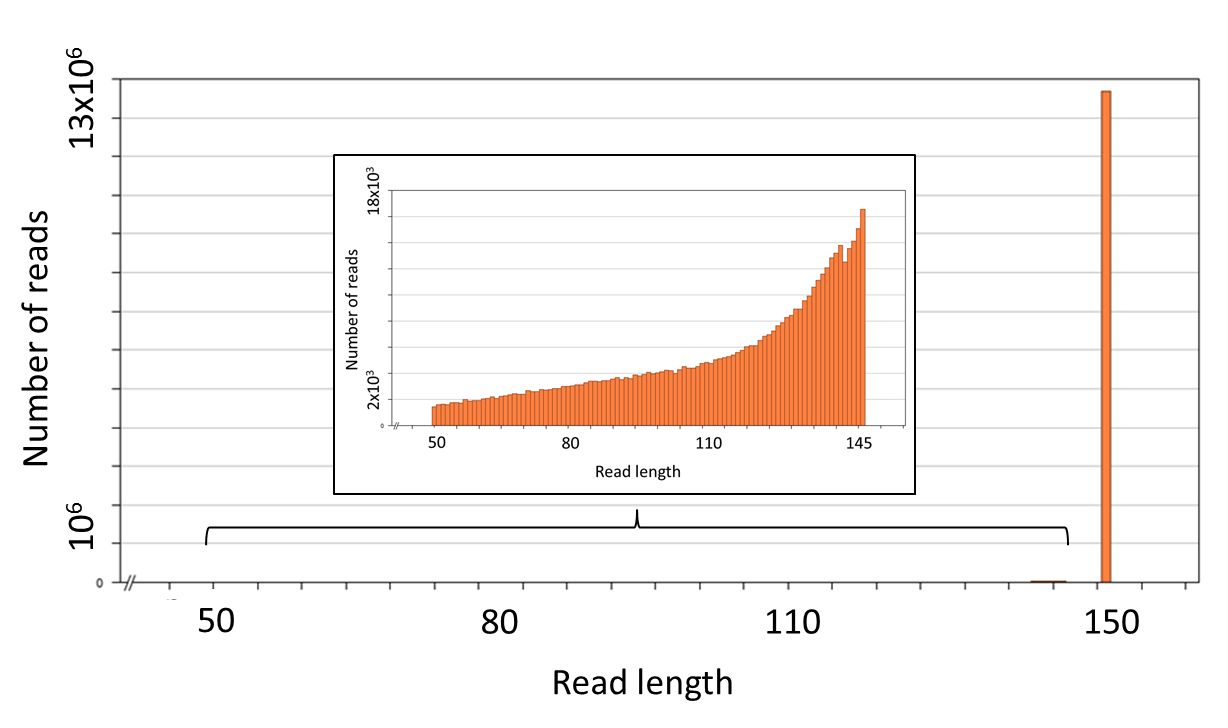

Fig. S1. Distribution of lengths of sequence reads of Myristica fragrans given in two graphs of different magnitudes. The outer graph corresponds to the number of reads in more than 95% of the sequences, all > 148 bp . The inner graph corresponds to the number of reads in the remaining sequences, all between 50 and 145 bp.

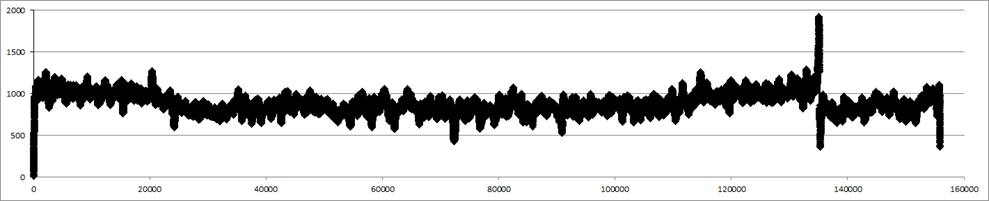

Fig. S2. Average coverage of base pair read position across the assembled genome of *Myristica fragrans*.
